# Time-dependent roles of adolescent- and adult-born dentate granule neurons in spatial learning

**DOI:** 10.1101/2020.05.08.084467

**Authors:** Nuria Masachs, Vanessa Charrier, Fanny Farrugia, Valerie Lemaire, Wilfrid Mazier, Nicolas Blin, Sophie Tronel, Marie-Françoise Montaron, Shaoyu Ge, Daniela Cota, Véronique Deroche-Gamonet, Cyril Herry, Djoher Nora Abrous

## Abstract

The dentate gyrus presents the peculiarity to be formed after birth in rodents. Adolescence is a very sensitive period during which cognitive competences are programmed. We investigated and compared the role of dentate neurons born during adolescence or generated during adulthood. We demonstrated that the ontogenetic stage of dentate neurons in relation to when they are generated dictates their participation in memory processes.

## INTRODUCTION

Adolescence is a critical period during which the brain undergoes a series of structural and functional maturation allowing the transition from childhood to adulthood^1, 2^. This period -ranging from approximately postnatal weeks 3-4 to 6-7 in rodents- is particularly important for the dentate gyrus (DG) growth, plasticity and emergence of social behavior^2^. The DG is also a key structure in learning and memory and appears as a heterogeneous structure composed by different populations of granule neurons (DGNs). Most of the DGNs in rodents are generated during the postnatal period, however, new granule neurons are added throughout life in a process called adult neurogenesis. It is believed that these DGNs generated during adulthood have a critical window during their development that confers them a unique role in hippocampal memory and once they are mature, they are functional indistinguishable to the other mature DGNs and they retire (for review see ^3^). In recent years, some investigations have focused on studying morphological and functional differences between adult DGNs and developmental DGNs. These studies have shown that these two populations present major morphological differences^4^; they undergo different survival processes^5^; that only adult DGNs but not developmental DGNs display experience-induced plasticity^6, 7^; and that they are recruited in different learning tasks^8^.

Although the role of dentate gyrus neurons born during adulthood (Adu-DGNs) in learning and memory has generated a great interest, the specific contribution of DGNs born during adolescence (dubbed here Ado-DGNs) in learning and memory is largely ignored. There are a few works where Ado-DGNs were manipulated, with stress^9, 10^, running^11^, social isolation^12^ mutation of the TLX gene^13, 14^, leading to behavioural consequences in adult animals on emotion and learning. However, new methods specifically targeting Ado-DGNs are needed in order to understand how they are implicated in learning and memory and if their role differ from Adu-DGNs’ one.

## RESULTS

To tackle this question, Ado-DGNs born in PN28 male rats were labeled using thymidine analogs (TA, **Table S1**) and their activation in response to spatial relational memory in the water maze (WM) was imaged using the expression of Zif268^8^, an immediate early gene (IEG) involved in the stabilization of long-lasting memories^15^. In a first experiment, rats were trained one month after the TA’s injection when animals have reached adulthood (**Figure1 a,b**). TA-labelled cells differentiated into neurons (% of TA expressing NeuN, Home Cage: 97.92±0.66%, Learning: 96.57±0.82%; t_43_=1.28, p=0.21) and were located in the inner granule cell layer (GCL). The percentage of activated Ado-DGNs (IdU-Zif268-IR cells) in the *Learning* group was superior to the one measured in the age-matched *Home Cage* control group (**Figure1 c,d**). Learning to find a visible platform (Cued learning), a hippocampal-independent task, or stress-physical activity (Swim group) did not activate Ado-DGNs (**Table S2** for statistical analysis). We then asked whether the age of these Ado-DGNs at the time of training was critical or whether they stably contributed to learning. We tagged Ado-DGNs and trained rats in the WM two or three months later (**Figure1 e,f,i,j**). Critically, under these conditions (2 or 3 months old Ado-DGNs), Ado-DGNs were not activated by training (**Figure1 g,h,k,l**) indicating that Ado-DGNs were activated by learning within a defined temporal window.

**Figure 1:**
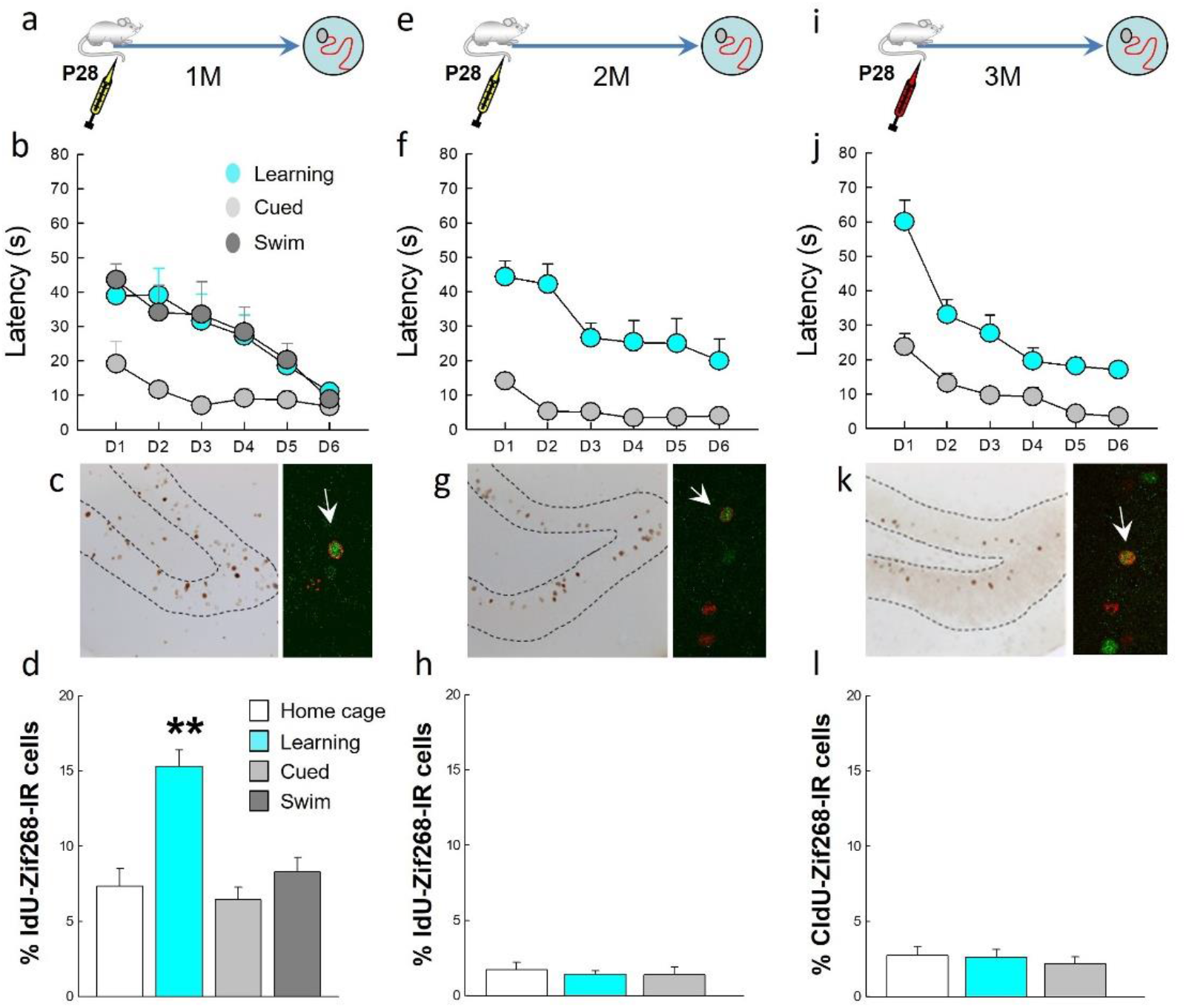
DGNs born in adolescents are recruited by spatial learning within a defined temporal window. (**a,e,j**) Experimental designs. (**b,f,j**) Learning curves. (**c,g,k**) Left panel: illustration of Idu- or CldU-labeled cells. Right panel: confocal example of Idu- or CldU-labeled cells Zif-268 (arrows indicate double labeled cells). Spatial learning increased the number of activated Adu-DGNs when the cells were 1 month old (**d**) and not when they were 2 months old (**h**) or 3 months old (**l**). ** p≤0.01 compared to other groups.

Then we asked whether the recruitment of Ado-DGNs was specific of this neuronal population by first focusing on neurons born during adulthood (Adu-DGNs, **Table S1, Figure S1a**). To this end, 2-month old rats were injected with BrdU and submitted to training one month later. We found that in this condition the percentage of BrdU-Zif268-IR neurons was not influenced by training (**Figure S1b, Table S2**). In contrast, when BrdU pulse and training interval was increased to 2 or 3 months, then a learning effect was observed (**Table S2, Figure S1b**). In contrast, neither 3-month old DGNs born during embryogenesis (Emb-DGNs) nor the neonatal period (Neo-DGNs) were activated by spatial learning in adult rats (**Table S1, Figure S1c,d, Table S2**). All together, these data suggest that : i) one month Ado-DGNs and *not* one month Adu-DGNs are recruited by learning and ii) Adu-DGNs (and not neurons born during development) are recruited when two and three month old.

To confirm the contribution of Ado-DGNs and Adu-DGNs in spatial learning, we used optogenetics to silence DGNs using a retrovirus expressing eGFP-tagged *Archaeorhodopsin* (RV-Arch-EGFP)^16^. We first validated this tool in rats by showing that optical stimulation reversibly silenced RV-Arch-EGFP-expressing neurons in a power-dependent manner as previously reported in mice^16^ (**Figure S2**). Then, we injected bilaterally the RV-Arch-EGFP into the DG of 28 days-old animals and two weeks before training, optic fibers were implanted bilaterally above the DG. Rats were trained in the WM 1, 2 or 4 months after tagging the cells. Half of the animals were trained with the light on (Ado-DGNs-Arch-Light) whereas the other half with the light off (Ado-DGNs-Arch-No-Light). No differences were observed between groups during the learning phase (**Figure S3a-c**). Seventy-two hours later, memory retention was measured by a probe test. During this test, the platform was removed and the optic fibers connected, but without light. When a delay of 1 month separated retroviral labeling from behavioral testing, both Ado-DGNs-Arch-No-light and Ado-DGNs-Arch-light animals remembered the platform position as they spend more time in the quadrant previously associated with the platform (Target quadrant, **Figure 2c, Table S3** for complete statistical analysis) compared to the other quadrants (O). In addition the time spent in the target quadrant was significantly different from the chance level (>25%). In subsequent experiments, rats were trained in the WM 2 or 4 months after tagging the cells. In contrast, retrieval was impaired in Ado-DGNs-Arch-light group when a delay of 2 months separated retroviral labeling from behavioral testing (**Figure 2f**; **Table S3** for complete statistical analysis). When the delay was increased to 4 months, optical inactivation did not alter memory retrieval (**Figure 2i**; **Table S2** for complete statistical analysis). To confirm that the effect of the light was not responsible of the impairment in the retrieval, an extra group of animals was injected bilaterally with RV-GFP without expressing *Archaeorhodopsin* and these animals were tested together with the other groups in the previous experiments (**Figure S4**). In none of the analyzed time points, we find an effect due to the light, neither in the learning (**Figure S4b,e,h**) nor in the retrieval (**Figure S4c,f,i**), confirming that the inhibition of a particular DGN population caused the incapacity of the animals to remember the platform location. After this verification, only the Arch-No-Light group was used as a control in the experiments.

**Figure 2:**
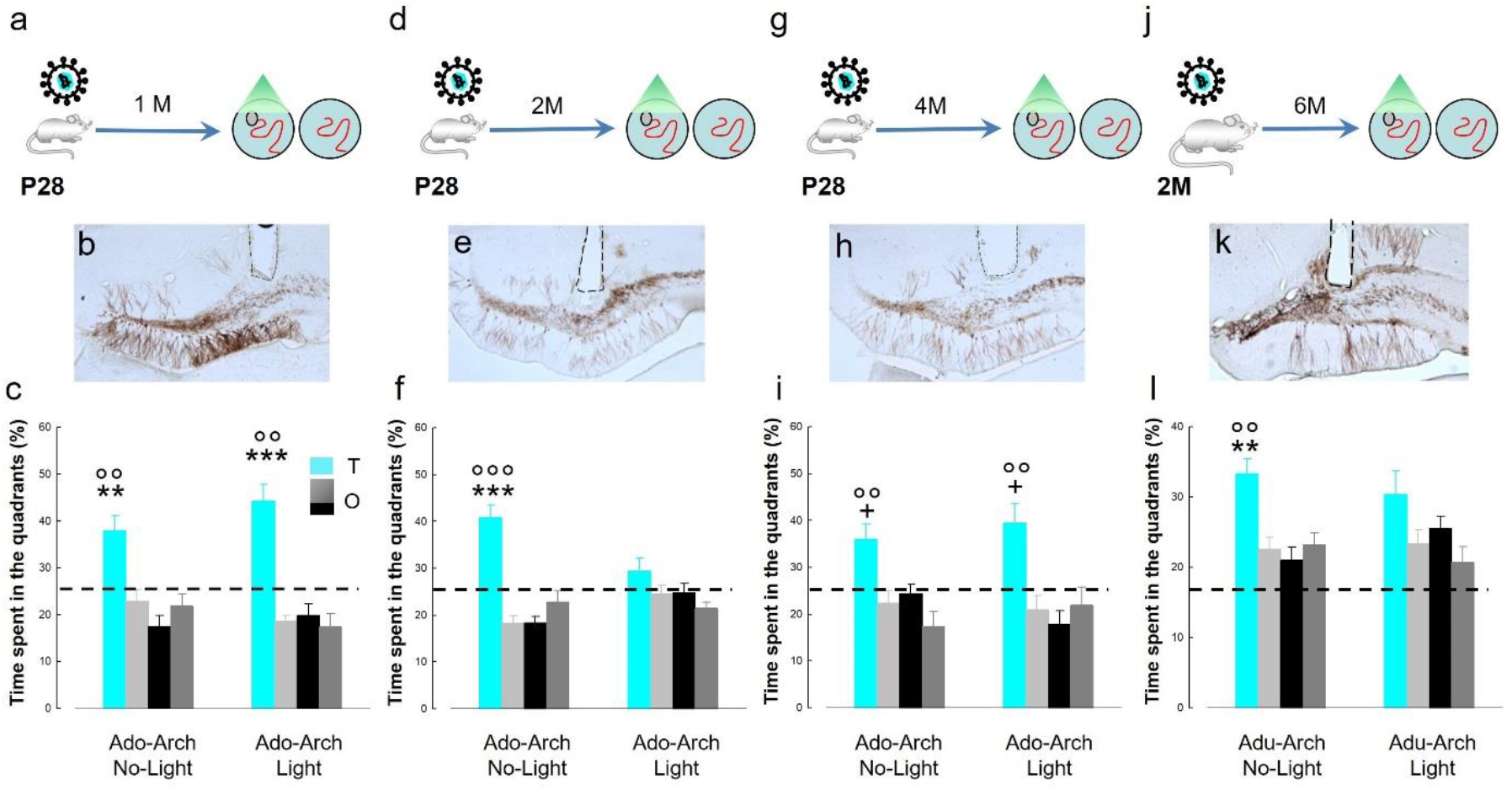
Optically silencing of Ado-DGNs and Adu-DGNs affects memory retrieval. (**a,d,g,j**) Experimental designs. Illustration of Ado-DGNs (**b,e,h**) or Adu-DGNs (**k**) infected with RV-Arch-EGFP; in dashed line, position where the optic fiber was placed. Silencing (**c**) 1-month old Ado-DGNs during training did not impair memory retention, whereas (**f**) silencing 2-months-old Ado-DGNs impaired the ability of the animals to remember the platform location. (**i**) When 4-months-old, silencing Ado-DGNs did not influence memory retention. (**l**) Silencing 6-month-old Adu-DGNs impaired the ability of the animals to remember the platform location. + at least at p<0.05 compared to other quadrants. **p≤0.01 ***p≤0.001 compared to other quadrants. °° p≤.01 and °°°p≤0.001 compared to chance level (25%). T: target quadrant. O: other quadrants.

To ascertain that the contribution of Ado-DGNs is due to the date of birth (Adolescence) *and* not the age (2 months) of the targeted population, RV-Arch-EGFP was injected bilaterally into the DG of 3 day-old animals that were trained 2 months later. Both Neo-DGNs-Arch-No-Light and Neo-DGNs-Arch-Light learned and remember the task (**Figure S5**).

Then we focused our attention on Adu-DGNs by injecting bilaterally RV-Arch-EGFP in 2-months-old animals and testing the impact of silencing Adu-DGNs 6 months later. This time point was chosen based on our previous experiments showing that Adu-DGNs are recruited by learning even when these neurons are six months old^8^. Half of the animals were trained with the light on (Adu-DGNs-Arch-Light) whereas the other half with the light off (Adu-DGNs-Arch-No-Light). No differences were observed between groups during the learning phase (**Figure S3d**). However, Adu-DGNs-Arch-Light animals were unable to remember the platform position at the probe test in contrast to their control group (**Figure 2l**; **Table S3** for complete statistical analysis). This result shows that when animals become older, memory retention depends on mature adult-born neurons.

Finally, the morphology of the dendritic arbor of DGNs born at different ontogenetic stage and the effect of light on their organization were analyzed. Clearly, neo-DGNs exhibit specific features compared to the other cohorts such as multiple primary dendrites, a wide branching angle and a more complex organization of the proximal part of the dendritic arbor (**Figure S6a-c, Figure3a**). When dendritic arbor of Ado-DGNs (aged of 1,2,4 months) was compared (**Figure S6b-d**), no major significant difference could be detected suggesting that their development is completed within few weeks. Adu-DGNs neurons (6-months of age) were, as previously described in mice, mainly similar to Ado-DGNs at the exception of the branch order that was higher (**Figure S6a-d, Figure3a**). Illumination had a different effect on dendritic length according to temporal origin of the DGNs (**Table S4**). It did not influence the dendritic organization of neo-DGNs (**Figure3b, Figure S6e**), consistently with the lack of an effect on memory. Similarly, Ado-DGNs dendritic length was not influenced by illumination (**Figure 3b, Figure S6f-h**). In contrast, the complexity of the dendritic arbor of mature Adu-DGNs was decreased by illumination, suggesting that these changes are responsible for impairment in retention (**Figure 3b-c**).

**Figure 3:**
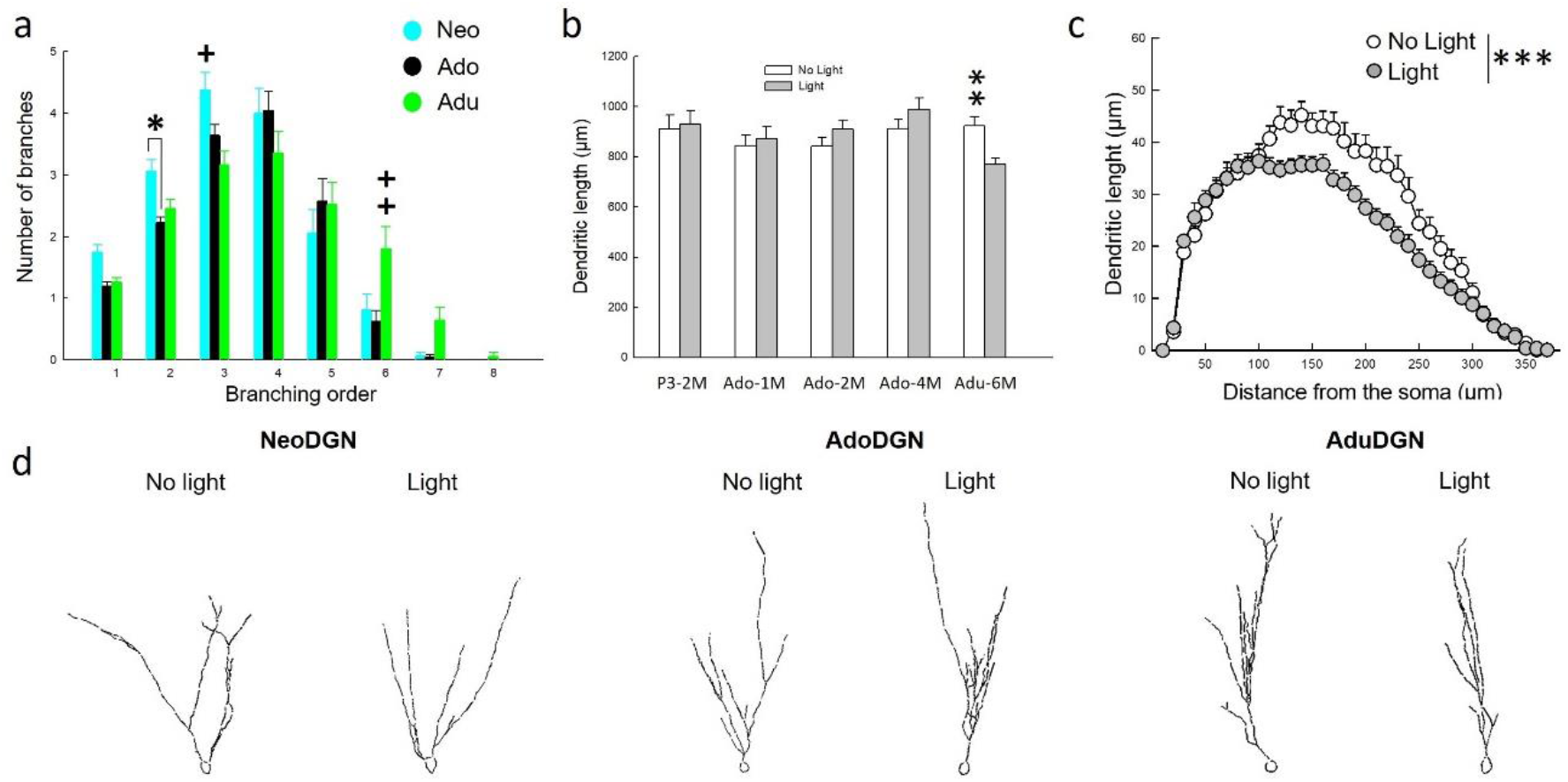
Different populations of DGNs show different branching arborization and they are distinctly modified by the light. **(a)** Dendritic branching shows divers profile between different DGNs population (interaction group x order : F_14,735_=3.630, p<0.0001), where Neo-DGNs show an increase in low order dendrites, while the Adu-DGNs an increase in higher order dendrites. Since no difference were observed between the populations in Ad-DGNs, only Ado-DGNs 4M is represented in the graph. **(b)** When the effect of light was examined on dendritic arbor, a reduction of the total length was only observed in Adu-DGNs, an effect **(c)** associated with a decrease in dendritic complexity (Table S4). **(d)** Representative tracings of the DGNs populations analyzed. * p≤0.05, **p≤0.01, ***p≤0.001, + at least at p<0.05 ++ at least at p≤0.01.

## DISCUSSION

Memory development is not a singular process and learning capacity emerges sequentially from the simple to the complex. The ability to learn spatial location develops later through adolescence into early adulthood. Four-to five-week-old animals use proximal cues to navigate (an independent-hippocampal strategy) and fail to search in the correct quadrant during probe trials^17, 18^. In contrast, 6 week-old animals use hippocampal-dependent strategy that requires the establishment of a cognitive map and the flexible use of the learned information (spatial relational memory). The maturation of spatial processing has been proposed to result from the maturation of the hippocampal formation and in particular of its spatially responsive cells^19^. Our results suggest that the emergence of spatial relational memory depends upon the recruitment of DGNs generated during adolescence.

Importantly and surprisingly this role appeared to be transient since Ado-DGNs were not anymore necessary when animals become older. This transient role was observed by imaging and silencing Ado-DGNs but the onset of the Ado-DGNs function was delayed when using optogenetic more probably because of the surgeries-induced inflammation that may delay neuronal development^20^. We previously reported that Neo-DGNs are recruited when animals have to discriminate different contexts and not to navigate through space^8^. Here we confirm that Neo-DGNs do not play a role in spatial memory processing. Given that when animals become adult, processing relational memory depends upon Adu-DGNs, we proposed that the adolescent wave of DGNs neurons might be involved in setting up the function (organizational value) whereas adult neurogenesis may rather have an adaptive value. In support of this contention, we have recently shown that Adu-DGNs -and not Ado-DGNs- are recruited by spatial learning when animals reached senescence ^21^.

This role of Ado-DGNs in shaping cognitive function and their specific role in affiliative behaviour^22^ may explain why adolescence compared to adulthood is very sensitive to the deleterious effects of chronic stress, drugs and high-fat diet^23-25^ on cognition and social behaviour.

In the search of the mechanism that might explain this different role, we focused on the dendritic structure of DGNs because the shape of a neuron’s dendritic tree influences how synaptic information is received and integrated, which can have functional implications^26^. In the DG in particular, it has been shown that dendritic architectures are predictive of GC activity during behavior spatial exploration, with DGNs in rats with high-order dendritic branching patterns being preferentially activated^27^.

Interestingly, we have identified different morphological features between the three mature populations: i) Neo-DGNs have more primary dendrites, a broader branching angle and more ramifications proximal to the soma compared to the two other populations, ii) Ado- and Adu-DGNs are indistinguishable in most of the parameters except that Adu-DGNs have a higher number of high order dendrites compare to the Ado-DGNs. Similarly in mice, we found that the dendritic organization of the DGNs depend upon the ontogenetic stage of the animals^4^. However, some species differences exist since in mice : i) mature Emb-DGNs and not Neo-DGNs exhibit more primary dendrites, ii) mature Neo-DGNs exhibit a longer dendritic length compared to Ado- and Adu-DGNs, iii) the number of high order dendrites in Ado-and Adu are similar^4^. Moreover, we could not observe a time-dependent evolution in Ado-DGNs’ morphology indicating that they reach the maturity within few weeks and mature faster than Adu-DGNs that keep on growing for at least 3 months (table S2 in ^7^).

When we compare the effect of inhibition in DGNs, we found a differential reactivity of mature DGNs born in the neonates, adolescent and adult animals. Developmentally born neurons were not sensitive to light inhibition in contrast to Adu-DGNs, indicating that Adu-DGNs are more plastic than developmental DGNs as we previously described^6, 7^. Even though we were expecting to see anatomical differences in Ado-DGNs 2 months to explain the impairment in retrieval, we did not find any effect of the light on the dendritic arbor. However, alterations in dendritic spines in pyramidal neurons following optogenetic stimulation have been previously reported^28, 29^ raising the possibility that light-induced alterations in dendritic spines of 2 month-old Ado-DGNs could be the responsible for the transient effect in the behavior. Further analysis is needed to verify this hypothesis.

In summary, our results highlight that DGNs born during adolescence and adulthood are two different populations with different neuronal features that share the same function but with a different timing and provide novel insights in disentangling how the heterogeneity of populations in the DG may contribute to differences in physiological and pathological states of hippocampal function.

## MATERIAL AND METHODS

### ANIMALS

A total of 219 male Sprague-Dawley rats (OFA, Janvier, France) were used (see Tables S1, S3). Rats were housed under a 12 h/12 h light/dark cycle (light off at 8pm) with *ad libitum* access to food and water. Temperature (22°C) and humidity (60%) were kept constant.

To study the immediate early genes activation in neurons born during adolescence, PN21 day-old rats were housed collectively until behavioral training. To study neurons born during development, pregnant female (n = 12, 3 months of age on delivery) Sprague-Dawley rats were individually housed in transparent cages. After birth, the litters were raised by their biological mothers until weaning (21 days after birth) and only the male progeny was kept after weaning. Rats were individually housed 1 week before the beginning of behavioral training.

For optogenetic experiments, 30 male rats of 6 weeks-old and 90 PN21 days-old rats on delivery were host in collective cages until the second surgery and moved afterwards in individual cages. For neonates DGN study, 43 male pups from 9 different females were used, they were raised by their biological mothers until weaning, and they were kept in collective cages until the second surgery. At the end of the experiments, only animals with GFP cells and ferules in good location were considered for further analysis (23, 64 and 20, respectively).

Experimental procedures have been carried out following the European directive of the parliament and the conceal of September 22, 2010 (2010/63/UE). Animal studies were approved by the ethical committee of the University of Bordeaux (CEEA50, Dir 13105, #5306).

### INJECTION OF THYMIDINE ANALOGS

BrdU (5-Bormo-2′-deoxyuridine), IdU (5-Iodo-2′-deoxyuridine) and CldU (5-chloro-2’-deoxyuridine) were dissolved in Phosphate Buffer (pH 8.4), 1N NH4OH/NaCl and NaCl, respectively. CldU and IdU injections were at equimolar doses of BrdU (50, 100 and 150 mg/kg (Table S1).

In the first three batches of animals, we examined whether neurons born during the preadolescent period were activated during spatial memory formation. Animals received one injection of IdU or CldU when 28 days-old. These animals were randomly distributed into 3 experimental groups (Home Cage, Cued and Learning) and were trained in the water maze (see below) one, two or three months after the injections.

In the fourth, fifth and sixth batches of animals, we examined whether neurons born during adulthood were activated during spatial memory. Animals received 3 injections of BrdU when 2 months-old. These animals were randomly distributed into 2 experimental groups (Home Cage and Learning) and were trained in the water maze one, two or three months after the injections.

In the seventh batch of animals, we examined whether neurons born during embryonic period and neurons born during the second postnatal week (PN14) in the same rats were activated during spatial memory formation. Pregnant female rats received two injections of CldU at E18.5 and E19.5. At 14 days-old (PN14), males received one injection of IdU. The animals were randomly distributed into 3 experimental groups (Home Cage, Cued and Learning) and trained in the water maze when 3 months-old.

### WATER MAZE TRAINING

The apparatus consisted of a circular plastic swimming pool (180 cm diameter, 60 cm height) in a room of 330 cm x 300 cm that was filled with water (20 ± 1 °C) rendered opaque by the addition of a white cosmetic adjuvant. Before training, the animals were habituated to the pool for 1 min per day for 2 days. During training, the Learning group was composed of animals that were required to locate a submerged platform, which was hidden 1.5 cm under the surface of the water in a fixed location, using the spatial cues available within the room (according to a previously described reference memory protocol ^30^). All rats were trained for four trials per day (90 s with an intertrial interval of 30 s and released from three starting points used in a pseudorandom sequence each day). If an animal failed to locate the platform, it was placed on that platform at the end of the trial. The time to reach the platform was recorded using a video camera that was secured to the ceiling of the room and connected to a computerized tracking system (Videotrack, Viewpoint). In the first 3 experiments, three control groups were used. The home cage group consisted of animals that were transferred to the testing room at the same time and with the same procedures as the Learning group but that were not exposed to the water maze. The Swim group was a Yoked control group for the stress and motor activity associated with the water maze training. The Swim group was composed of rats that were placed into the pool without the platform and were paired for the duration of the trial with the Learning animals. The third control group was the Cued group, which was composed of rats that were trained to find a visible platform in a fixed location. Animals in this group were tested for 4 trials per day (90 s with an intertrial interval of 30 s and beginning from three different starting points used in a pseudorandom sequence). Trained animals and different age-matched control groups were perfused transcardially 90 min after the last trial.

### RETROVIRUS PRODUCTION

The murine Moloney leukemia virus-based retroviral vector CAG-GFP has been previously described ^6^ and is a gift from D r FH Gage (Salk Institute, La Jolla, CA). The retrovirus expressing an inhibitory optogene *Archaerhodopsin*-3 (Arch-EGFP) was generated by Dr Ge S^16^. High titers of retroviruses (between 5×10^8^ and 5×10^9)^ were prepared with a human 293-derived retroviral packaging cell line (293GPG)^31^ kindly provided by Dr Dieter Chichung Lie (Friedrich-Alexander Universität Erlangen-Nürnberg, Erlangen, Germany). Virus-containing supernatant was harvested three days after transfection with Lipofectamine 2000 (Thermofisher). This supernatant was then cleared from cell debris by centrifugation at 3500 rpm for 15 min and filtered through a 0.45 µm filter (Millipore). Viruses were concentrated by two rounds of centrifugation (19 500 rpm 2h) and resuspended in PBS.

### STEROTAXIC INJECTIONS OF RETROVIRUS

The retroviral solution was injected bilaterally onto the septal part of the DG. Rats were anesthetized in a box using a mix of oxygen and 3% isoflurane and maintained asleep with 2% isoflurane during the whole surgery in a stereotaxic apparatus. Animals also received lidocaine (0.1 ml, sub-cutaneous sc, Vetoquinol) on the incision site and metacam (1ml/kg, sc, Boehringer Ingelheim). Glass micropipettes connected to a Hamilton syringe placed into a micro injector (KDS legato 130) directly attached to the stereotaxic apparatus were used. Injections were realized respecting the following coordinates^32^: **For P3 groups**, [AP]: - 1.2 mm; lateral [L]: ± 1.35 mm; deepness [P]: - 2.6 mm; **P28 groups**, [AP]: −2.8 mm; lateral [L]: ±1.5 mm; deepness [P]: −3.5 mm and for **Adult groups**, [AP]: −3.3 mm; lateral [L]: ±1.6 mm; deepness [P]: −4.2 mm. One (P3), two (P28) or three µl (Adult) of viral suspension were injected at a flow rate 0.25L/min (8min). Cannulas were maintained in position for 3 minutes after the end of the injections to let the suspension infuse. Then the skin was stitched with absorbable sutures and the rat was placed in a recovery chamber (37°C) until it woke up.

### ELECTROPHYSIOLOGICAL RECORDING

Animals were deeply anesthetized (Xylazine 16.7mg/kg plus Ketamine 167mg/kg, intraperitoneal, Bayer santé and Mérial, Germany respectively) and sacrificed. Dissected brain was immediately immerged in ice-cold oxygenated cutting solution (in mM: 180 Sucrose, 26 NaHCO3, 11 Glucose, 2.5 KCl, 1.25 NaH2PO4, 12 MgSO4, 0.2 CaCl2, saturated with 95% O2-5% CO2). 350 mm slices were obtained using a vibratome (VT1200S Leica, Germany) and transferred into a 34°C bath of oxygenated aCSF (in mM: 123 NaCl, 26 NaHCO3, 11 Glucose, 2.5 KCl, 1.25 NaH2PO4, 1.3 MgSO4, 2.5 CaCl2; osmolarity 310 mOsm/l, pH 7.4) for 30 minutes and then cooled down progressively till room temperature (RT; 23-25°C) in oxygenated aCSF. After a 45 min recovery period at RT, slices were anchored with platinum wire at the bottom of the recording chamber and continuously bathed in oxygenated aCSF (RT; 2ml/min) during recording.

Fluorescent DGNs cells were identified using fluorescence/infrared light (pE-2 CoolLED excitation system, UK). Neurons action potential firing was monitored in whole-cell current-clamp recording configuration. Patch electrodes were pulled (micropipette puller P-97, Sutter instrument, USA) from borosilicate glass (O.D. 1.5 mm, I.D. 0.86 mm, Sutter Instrument) to a resistance of 2-4 mΩ. The pipette internal solution contained [in mM: 125 potassium gluconate, 5 KCl, 10 Hepes, 0.6 EGTA, 2 MgCl2, 7 Phosphocreatine, 3 adenosine-5’-triphosphate (magnesium salt), 0.3 guanosine-5’-triphosphate (sodium salt) (pH adjusted to 7.25 with KOH; osmolarity 300 mOsm/l adjusted with d-Mannitol)] and added with biocytin 0.4% (liquid junction potential - 14.8mV was corrected on the data and statistics). Electrophysiological data were recorded using a Multiclamp 700B amplifier (Molecular devices, UK), low-pass filtered at 4 kHz and digitized at 10Hz (current clamp) or 4 Hz (voltage clamp) (Digidata 1440A, Molecular devices, UK). Activation of Arch was obtained after illumination of the recorded area at 570 nm (pE-2 CoolLED excitation system, UK). Signals were analyzed offline (Clampfit software, pClamp 10, Molecular devices, UK).

### OPTIC FIBER IMPLANTATION

Home-made optic fibers were fixed in the ferules. The implantation step was realized during a second surgery following the same protocol describe before. The skull was scratched and three anchors screws were fixed (one on the anterior part and two on the posterior part). The two optic fibers (length: 4.5 mm, diameter: 200 µm, numeric aperture (NA): 0.37) were lowered with a speed of ∼ 2mm/min upward to the RV injection site respecting the following stereotaxic coordinates^32^: **Adult group**: [AP]: −3.3 mm; [L]: ±1.6 mm; [P]: −4 mm from skull surface; **Adolescence group 1 month delay:** [AP]: −3.2 mm; [L]: ±1.5 mm; [P]: −3.6mm from bregma; **Adolescence group 2 and 4 months delay:** [AP]: −3.4 mm; [L]: ±1.5 mm; [P]: −3.8; **Neonatal group:** [AP]: −3.3 mm; [L]: ±1.6 mm; [P]: −4. Fibers were stabilized using dental cement. When the cement was dry, scalps were sutured and disinfected with local antiseptic treatment (Betadine). All rats were followed up for until the starting of the behavior. During the handling, fur was checked and their scar disinfected systematically. Rats were handled every day to habit them for the behavioral procedure.

### OPTOGENETIC MANIPULATION OF DGNS DURING TRAINING IN THE WATER MAZE

The water maze (circular pool with a diameter of 1.80m and 60cm deep) was placed in a room (375 cm x 370 cm) equipped with a green laser source (OptoDuet 532,5nm, 200mW, Ikecool) that was connected to a rotatory joint to avoid tangling of the patch cords connected to the intra-dentate optic fibers. Light intensity at the optic fiber cable tip was controlled every day before the beginning of the session (8-10 mV). Illumination was applied during the whole session at 1Hz pulses of 750ms. Same protocol described before was used for the training. The time to reach the platform was recorded using a video camera connected to a computerized tracking system (Polyfiles, Imetronic). When animals reached the plateau phase, training was stopped and 72h later animals were tested to find the platform. During the probe test, the hidden platform was removed and the time spent in each quadrant was quantified.

### IMMUNOHISTOCHEMISTRY

Animals were perfused transcardially with a phosphate buffered solution of 4 % paraformaldehyde. After 1 week of fixation, brains were cut using a vibratome as previously described ^30^. Free-floating sections (50 µm) were processed using a standard immunohistochemical procedure to visualize the thymidine analogs (BrdU, CldU, IdU) and retrovirus expressing EGFP in alternating one-in ten sections using different anti-BrdU antibodies from different vendors (for BrdU: mouse primary at 1/100, Dako; CldU: rat primary at 1/1,000, Accurate Chemical; IdU: mouse primary at 1/1,000, BD Biosciences) and with a rabbit anti GFP (Millipore, 1:2000). Bound antibodies were visualized with horse anti-mouse (1/200, Abcys for BrdU and Idu), goat anti-rat (1/200, Vector for CldU) and goat anti-rabbit (1:200). The number of XdU-immunoreactive (IR) cells in the granule and subgranular layers (GCL) of the DG was estimated on a systematic random sampling of every tenth section along the septo-temporal axis of the hippocampal formation using a modified version of the optical fractionator method. Indeed, all of the X-IR cells were counted on each section and the resulting numbers were tallied and multiplied by the inverse of the section sampling fraction (1/ssf=10).

The activation of embryonic, neonatale, adolescent and adult generated neurons was examined using immunohistofluorescence. To visualize the cells that incorporated thymidine analogs, one section out of ten was incubated with BrdU antibodies (BrdU, 1/1,000, CldU, 1/500, Accurate Chemical; IdU, 1/500, BD Bioscience). Bound antibodies were visualized with Cy3-goat anti-rat antibodies (1/1,000, Jackson for CldU and BrdU) or Cy3-goat anti-mouse antibodies (1/1,000, Jackson for IdU). Sections were also incubated with rabbit anti-Zif268 (1/500, Santa Cruz Biotechnology). Bound antibodies were visualized with Alexa488-goat anti-rabbit antibodies (1/1,000, Invitrogen). Primary antibodies for CldU (or IdU or BrdU) and IEG (Zif268) were incubated simultaneously at 4 °C for 72 h, and secondary antibodies were incubated simultaneously at RT for 2 h. CldU-IEG and IdU-IEG labeling were not analyzed on the same sections, given the cross-reactivity between the secondary (and not primary) antibodies (Tronel et al., 2015). The sections were then incubated with the two secondary antibodies. The phenotype of the CldU-IR cells was examined by immunofluorescence labeling using NeuN (1:1,000; Millipore) revealed with a Cy5-goat anti-mouse (1:1,000; Jackson) and the phenotype of IdU-IR cells using Calbindin (Calb, 1:250; Santa Cruz Biotechnologies) revealed with Alexa-Fluor 647 donkey anti-goat (1/1000; Jackson). Double labeling was determined using a SPE confocal system with a plane apochromatic 63X oil lens (numerical aperture 1.4; Leica). The percentage of XdU cells expressing IEG and the number of XdU-cells expressing NeuN or Calb were calculated as follows: (Nb of XdU^+^/IEG^+^ cells)/[(Nb of XdU^+^/IEG^-^ cells) + (Nb of XdU^+^/IEG^+^ cells)] x 100. All sections were optically sliced in the Z plane using 1 µm interval and cells were rotated in orthogonal planes to verify double labelling. In all experiments, a minimum of 200 developmentally generated neurons were analyzed per rat.

The morphometric analysis of virus-labeled neurons was performed with a x100 objective as previously described ^4^. Measurements of dendritic parameters as well as sholl analysis were performed with the Neurolucida (software Microbrightfield, Colechester, VT, USA). Only neurons from suprapyramidal layer were analyzed. Data shown is from at least 5 rats per group with a minimum of 4 neurons per group (**Table S4**).

### STATISTICAL ANALYSIS

The data (mean ± s.e.m.) were analyzed using t-test and ANOVA, which was followed by the Newman-Keuls test when necessary.

## ACKNOWLEDGEMENTS

The authors thank Dr F Massa and Dr G Marsicano for lending their electrophysiology equipment. We greatly acknowledge C Dupuy for animal care. Supported by Inserm (to DNA), FRM (to DNA), LabEx BRAIN ANR-10-LABX-43 (Bordeaux Region Aquitaine Initiative for Neuroscience to DNA), ANR [grant ANR-10-EQX-008-1 to VDG, CH, DC, DNA, EquipEx OptoPath™] ANR Neuronutrisens ANR-13-BSV4-0006 (to DC). NM was a recipient of a post-doctoral study grant from “La Fondation FYSSEN.” This work benefited from the support of the Biochemistry and Biophysics Facility of the Bordeaux Neurocampus funded by the LabEX BRAIN ANR-10-LABX-43 and the Animal Housing facility funded by Inserm and LabEX BRAIN ANR-10-LABX-43. The confocal analysis was done in the Bordeaux Imaging Center (BIC), a service unit of the CNRS-INSERM and Bordeaux University, member of the national infrastructure France BioImaging supported by the French National Research Agency (ANR-10-INBS-04).

## Abbreviations

BrdU: 5-bromo-2’-deoxyuridine;
CldU: 5-chloro-2’-deoxyuridine;
DG: dentate gyrus;
IdU: 5-iodo-2’-deoxyuridine.
DGN: dentate granule neurons,
Adu: Adult,
Ado: Adolescent,
Emb: embryonic,
Neo: Neonate.
TA: thymidine analog.

## Reference List

1. Fuhrmann, D., Knoll, L.J., & Blakemore, S.J. Adolescence as a Sensitive Period of Brain Development. Trends Cogn Sci. 19, 558–566 (2015).

2. Spear, L.P. The adolescent brain and age-related behavioral manifestations. Neurosci. Biobehav. Rev. 24, 417–463 (2000).

3. Abrous, D.N. & Wojtowicz, J.M. Interaction between Neurogenesis and Hippocampal Memory System: New Vistas. Cold Spring Harb. Perspect. Biol. 7, (2015).

4. Kerloch, T., Clavreul, S., Goron, A., Abrous, D.N., & Pacary, E. Dentate Granule Neurons Generated During Perinatal Life Display Distinct Morphological Features Compared With Later-Born Neurons in the Mouse Hippocampus. Cereb. Cortex 29, 3527–3539 (2019).

5. Cahill, S.P., Yu, R.Q., Green, D., Todorova, E.V., & Snyder, J.S. Early survival and delayed death of developmentally-born dentate gyrus neurons. Hippocampus(2017).

6. Lemaire, V. et al. Long-lasting plasticity of hippocampal adult-born neurons. J Neurosci 32, 3101–3108 (2012).

7. Tronel, S. et al. Spatial learning sculpts the dendritic arbor of adult-born hippocampal neurons. Proc Natl Acad Sci U. S. A 107, 7963–7968 (2010).

8. Tronel, S., Lemaire, V., Charrier, V., Montaron, M.F., & Abrous, D.N. Influence of ontogenetic age on the role of dentate granule neurons. Brain Struct. Funct. 220, 645–661 (2015).

9. Hueston, C.M., Cryan, J.F., & Nolan, Y.M. Stress and adolescent hippocampal neurogenesis: diet and exercise as cognitive modulators. Transl. Psychiatry 7, e1081 (2017).

10. McCormick, C.M. et al. Social instability stress in adolescent male rats alters hippocampal neurogenesis and produces deficits in spatial location memory in adulthood. Hippocampus 22, 1300–1312 (2012).

11. O’Leary, J.D. et al. Differential effects of adolescent and adult-initiated exercise on cognition and hippocampal neurogenesis. Hippocampus 29, 352–365 (2019).

12. Hueston, C.M., Cryan, J.F., & Nolan, Y.M. Adolescent social isolation stress unmasks the combined effects of adolescent exercise and adult inflammation on hippocampal neurogenesis and behavior. Neuroscience 365, 226–236 (2017).

13. Kozareva, D.A., O’Leary, O.F., Cryan, J.F., & Nolan, Y.M. Deletion of TLX and social isolation impairs exercise-induced neurogenesis in the adolescent hippocampus. Hippocampus 28, 3–11 (2018).

14. Kozareva, D.A., Foley, T., Moloney, G.M., Cryan, J.F., & Nolan, Y.M. TLX knockdown in the dorsal dentate gyrus of juvenile rats differentially affects adolescent and adult behaviour. Behav. Brain Res. 360, 36–50 (2019).

15. Veyrac, A., Besnard, A., Caboche, J., Davis, S., & Laroche, S. The transcription factor Zif268/Egr1, brain plasticity, and memory. Prog. Mol. Biol. Transl. Sci. 122, 89–129 (2014).

16. Gu, Y. et al. Optical controlling reveals time-dependent roles for adult-born dentate granule cells. Nat. Neurosci 15, 1700–1706 (2012).

17. Schenk, F. Development of place navigation in rats from weaning to puberty. Behav. Neural Biol 43, 69–85 (1985).

18. Rudy, J.W., Stadler-Morris, S., & Albert, P. Ontogeny of spatial navigation behaviors in the rat: dissociation of “proximal”- and “distal”-cue-based behaviors. Behav. Neurosci. 101, 62–73 (1987).

19. Langston, R.F. et al. Development of the spatial representation system in the rat. Science 328, 1576–1580 (2010).

20. Monje, M.L., Toda, H., & Palmer, T.D. Inflammatory blockade restores adult hippocampal neurogenesis. Science 302, 1760–1765 (2003).

21. Montaron, M.F., Charrier, V., Blin, N., Garcia, P., & Abrous D.N. Responsiveness of dentate neurons generated throughout adult life determines spatial memory ability of aged rats. Aging Cell. 2020. Ref Type: In Press

22. Wei, L., Meaney, M.J., Duman, R.S., & Kaffman, A. Affiliative behavior requires juvenile, but not adult neurogenesis. J Neurosci 31, 14335–14345 (2011).

23. Spear, L.P. Heightened stress responsivity and emotional reactivity during pubertal maturation: Implications for psychopathology. Dev. Psychopathol. 21, 87–97 (2009).

24. Boitard, C. et al. Juvenile, but not adult exposure to high-fat diet impairs relational memory and hippocampal neurogenesis in mice. Hippocampus 22, 2095–2100 (2012).

25. Boitard, C. et al. Switching Adolescent High-Fat Diet to Adult Control Diet Restores Neurocognitive Alterations. Front Behav. Neurosci. 10, 225 (2016).

26. Lefebvre, J.L., Sanes, J.R., & Kay, J.N. Development of dendritic form and function. Annu. Rev. Cell Dev. Biol. 31, 741–777 (2015).

27. Diamantaki, M., Frey, M., Berens, P., Preston-Ferrer, P., & Burgalossi, A. Sparse activity of identified dentate granule cells during spatial exploration. Elife. 5, (2016).

28. Mendez, P., Stefanelli, T., Flores, C.E., Muller, D., & Luscher, C. Homeostatic Plasticity in the Hippocampus Facilitates Memory Extinction. Cell Rep. 22, 1451–1461 (2018).

29. Moulin, T.C. et al. Chronic in vivo optogenetic stimulation modulates neuronal excitability, spine morphology, and Hebbian plasticity in the mouse hippocampus. Hippocampus 29, 755–761 (2019).

30. Dupret, D. et al. Spatial learning depends on both the addition and removal of new hippocampal neurons. PLOS Biology 5, 1683–1694 (2007).

31. Ory, D.S., Neugeboren, B.A., & Mulligan, R.C. A stable human-derived packaging cell line for production of high titer retrovirus/vesicular stomatitis virus G pseudotypes. Proc. Natl. Acad. Sci. U. S. A 93, 11400–11406 (1996).

32. Paxinos, G. & Watson, C. The rat brain in sterotaxic coordinates (Academic Press, 1982).

