## Supplementary material for "Time-dependent roles of adolescent- and adult-born dentate granule neurons in spatial learning"

| Batches | Experiments | Animal's age at the time of XdU injection | Animals 'age at the time of sacrifice | Cells' age at the time of training | XdU dose (mg/kg) | Group size |
|---|---|---|---|---|---|---|
| **Ado**-DGNs | Activation of neurons generated during adolescent period | 28 days | 2 months | 1 month | IdU : 1x150 | Control = 9<br>Learning = 11<br>Swim= 9<br>Cued = 9 |
| **Ado**-DGNs | Activation of neurons generated during adolescent period | 28 days | 3 months | 2 months | IdU: 1x100 | Control = 5<br>Learning = 9<br>Cued = 5 |
| **Ado**-DGNs | Activation of neurons generated during adolescent period | 28 days | 4 months | 3 months | CldU: 1x50 | Control = 5<br>Learning = 6<br>Cued = 6 |
| **Adu**-DGNs | Activation of neurons generated during adulthood | 2 months | 3 months | 1 month | BrdU: 3 x 100 | Control = 3<br>Learning = 6 |
| **Adu**-DGNs | Activation of neurons generated during adulthood | 2 months | 4 months | 2 months | BrdU: 3 x 100 | Control = 3<br>Learning = 5 |
| **Adu**-DGNs | Activation of neurons generated during adulthood | 2 months | 5 months | 3 months | BrdU: 3 x 100 | Control = 5<br>Learning = 7 |
| **Emb**-DGNs<br>**Neo**-DGNs | Activation of neurons generated during embryonic and neonatal periods | E18.5<br>14 days | 3 months | 10 weeks<br>12 weeks | CldU: 2x50<br>IdU: 1x50 | Control = 6<br>Learning = 13<br>Cued = 8 |

**Table S1: Summary of the experimental groups and experiments**

| Experiments | Delay | Nb of XdU-IR cells | Comparison of the number of XdU–IR cells | % of XdU-Zif-268IR cells | Comparison of activated DGNs-IR cells |
|---|---|---|---|---|---|
| **Ado**-DGNs | 1 M | Control : 18322 ±617<br>Learning : 20351 ± 1384<br>Cued : 23040 ± 1489<br>Swim : 22904 ± 1688 | $F_{3,37}$=2.61; p=0.07 | Control : 7.33 ± 1.2<br>Learning : 15.30 ± 1.09<br>Cued : 8.29 ± 0.95<br>Swim : 6.45 ± 0.80 | $F_{3,37}$=16.67 ; p=0.001 |
| **Ado**-DGNs | 2 M | Control : 18948 ± 2848<br>Learning : 15980 ± 1257<br>Cued: 16140 ± 940 | $F_{2,18}$=0.85; p=0.45 | Control : 1.70 ± 0.50<br>Learning : 1.41 ± 0.23<br>Cued: 1.37 ± 0.51 | $F_{2,18}$=0.20, p=0.82 |
| **Ado**-DGNs | 3 M | Control : 11068 ± 1752<br>Learning : 11680 ± 783<br>Cued : 10700 ± 975 | $F_{2,16}$=0.20; p=0.83 | Control : 2.75 ± 0.59<br>Learning 2.59 ± 0.55<br>Cued : 2.17 ± 0.47 | $F_{2,16}$=0.32, p=0.74 |
| **Adu**-DGNs | 1 M | Control : 10013 ± 2713<br>Learning : 9583 ± 1212 | $t_7$= 0.17; p=0.86 | Control : 3.87 ± 1.36<br>Learning : 3.83 ± 0.73 | $t_7$= 0.03; p=0.98 |
| **Adu**-DGNs | 2 M | Control : 9360 ± 1655<br>Learning : 11240 ± 887 | $t_6$= -1.11; p=0.31 | Control : 6.67 ± 2.79<br>Learning : 17.69 ± 1.50 | $t_6$= -3.86; p=0.008 |
| **Adu**-DGNs | 3 M | Control : 5792 ± 1038<br>Learning : 5580 ± 717 | $t_{10}$= 0.17; p=0.86 | Control : 11.75 ± 0.74<br>Learning : 19.28 ± 1.55 | $t_{10}$= -3.84; p=0.003 |
| **Emb**-DGNs | 3 M | Control : 132508 ± 9219<br>Learning : 152655± 14733<br>Cued : 136854 ± 14310 | $F_{2,25}$= 0.50; p=0.61 | Control : 5.34 ± 1.12<br>Learning : 6.44 ± 0.73<br>Cued : 5.14 ± 0.55 | $F_{2,25}$=0.92, p=0.41 |
| **Neo**-DGNs | 10 W | Control : 19407 ± 2115<br>Learning : 23476 ± 1268<br>Cued : 21663 ± 1262 | $F_{2,25}$=0.77; p=0.19 | Control : 8.72 ± 1.30<br>Learning : 8.03 ± 0.99<br>Cued : 7.83 ± 0.87 | $F_{2,25}$=0.14, p=0.87 |

**Table S2: Statistical analysis of the expression of Zif268 in DGNs.**

| DGNs population | Delay | Group size | Number of GFP-IR cells | Analysis of GFP-IR cell numbers | Analysis of time spent in the quadrants | Comparison of TQ to chance level |
|---|---|---|---|---|---|---|
| **Ado**-DGNs-Arch-No-Light | 1 M | 14 | 2454 ± 400 | $F_{2,29}$=0.37, p=0.69 | $F_{3,39}$=8.59, p<0.001 with TQ<SE p=0.0013; TQ<NE p<0.001; TQ<SW p=0.0016. | $t_{13}$=4.07, p=0.0013 |
| **Ado**-DGNs-Arch-Light | | 6 | 2395 ± 418 | | $F_{3,15}$=17.23, p<0.001 with TQ< SE p<0.001; TQ<NE p<0.001; TQ<SW p<0.001 | $t_{5}$=5.35, p=0.0031 |
| **Ado**-DGNs-GFP-Light | | 12 | ± 418 | | $F_{3,33}$=10.29, p<0.001 with TQ< SE p<0.001; TQ<NE p<0.001; TQ<SW p<0.001 | $t_{11}$=3.89, p=0.003 |
| **Ado**-DGNs-Arch-No-Light | 2 M | 13 | 3498 ± 670 | $F_{2,36}$=2.45, p=0.10 | $F_{3,36}$=19.9, p<0.001 with TQ<SE p<0.001; TQ<NE p<0.001; TQ<SW p<0.001 | $t_{12}$=5.86, p<0.001 |
| **Ado**-DGNs-Arch-Light | | 13 | 4849 ± 541 | | $F_{3,36}$=1.84, p=0.16 | $t_{12}$=1.53, p=0.15 |
| **Ado**-DGNs-GFP-Light | | 13 | 2925 ± 675 | | $F_{3,36}$=15.12, p<0.001 with TQ< SE p<0.001; TQ<NE p<0.001; TQ<SW p<0.001 | $t_{12}$=5.17, p<0.001 |
| **Ado**-DGNs-Arch-No-Light | 4 M | 10 | 2801 ± 634 | $F_{2,24}$=1.55, p=0.23 | $F_{3,27}$=5.8, p=0.003 with TQ<SE p=0.0176; TQ<NE p=0.0185; TQ<SW p=0.0024 | $t_{9}$=3.29, p=0.009 |
| **Ado**-DGNs-Arch-Light | | 8 | 3885 ± 457 | | $F_{3,21}$=5.6, p=0.005 with TQ<SE p=0.013; TQ<NE p=0.0068; TQ<SW p=0.0068 | $t_{7}$=3.40, p=0.01 |
| **Ado**-DGNs-GFP-Light | | 9 | 4151 ± 617 | | $F_{3,24}$=14.85, p<0.001 with TQ< SE p<0.001; TQ<NE p<0.001; TQ<SW p<0.001 | $t_{8}$=5.06, p=0.001 |
| **Adu**-DGNs-Arch-No-Light | 6 M | 11 | 2129 ± 365 | $t_{21}$=1.49, p=0.15 | $F_{3,30}$=6.4, p=0.001 with TQ<SE p=0.005; TQ<NE p=0.002; TQ<SW p=0.003 | $t_{10}$=3.82, p=0.003 |
| **Adu**-DGNs-Arch-Light | | 12 | 2929 ± 389 | | $F_{3,33}$=2.2, p=0.1 | $t_{11}$=1.62, p=0.13 |
| **Neo-DGNs** Arch-No-Light | 2 M | 7 | 6117 ± 2876 | $t_{18}$=1.01, p=0.33 | $F_{3,18}$=12.24, p<0.001 with TQ<SE p<0.001; TQ<NE p<0.001; TQ<SW p<0.001 | $t_{6}$=5.18, p=0.002 |
| **Neo-DGNs** Arch—Light | | 13 | 9126 ± 1561 | | $F_{3,36}$=6.18, p=0.0017 with TQ<SE p=0.002; TQ<NE p=0.003; TQ<SW p=0.021 | $t_{12}$=3.23, p=0.0072 |

**Table S3: Statistical analysis of behaviour performance in optogenetic experiments.**

| Experiments | Delay | Number of rats | Number of DGNs analyzed | Light effect in Sholl analysis | Light effect in total dendritic length |
|---|---|---|---|---|---|
| **Ado**-DGNs-Arch-No-Light | 1 M | 6 | 34 | $F_{1,68}$=0.1490, p=0.7007 | $t_{68}$=0.3878, p=0.6994 |
| **Ado**-DGNs-Arch-Light | | 6 | 36 | | |
| **Ado**-DGNs-Arch-No-Light | 2 M | 10 | 59 | $F_{1,125}$=1.752, p=0.1880 | $t_{125}$=1.325, p=0.1876 |
| **Ado**-DGNs-Arch-Light | | 13 | 68 | | |
| **Ado**-DGNs-Arch-No-Light | 4 M | 8 | 45 | $F_{1,85}$=1.807, p=0.1824 | $t_{85}$=1.344, p=0.1824 |
| **Ado**-DGNs-Arch-Light | | 8 | 42 | | |
| **Adu**-DGNs-Arch-No-Light | 6 M | 6 | 31 | $F_{1,83}$=12.96, p=0.0005 | $t_{83}$=3.603, p=0.0005 |
| **Adu**-DGNs-Arch-Light | | 11 | 54 | | |
| **Neo-DGNs** Arch-No-Light | 2 M | 5 | 32 | $F_{1,60}$=0.011, p=0.9136 | $t_{60}$=0.1084, p=0.9140 |
| **Neo-DGNs** Arch-Light | | 5 | 30 | | |

**Table S4: Statistical analysis of light effect in dendritic morphology of DGNs populations.**

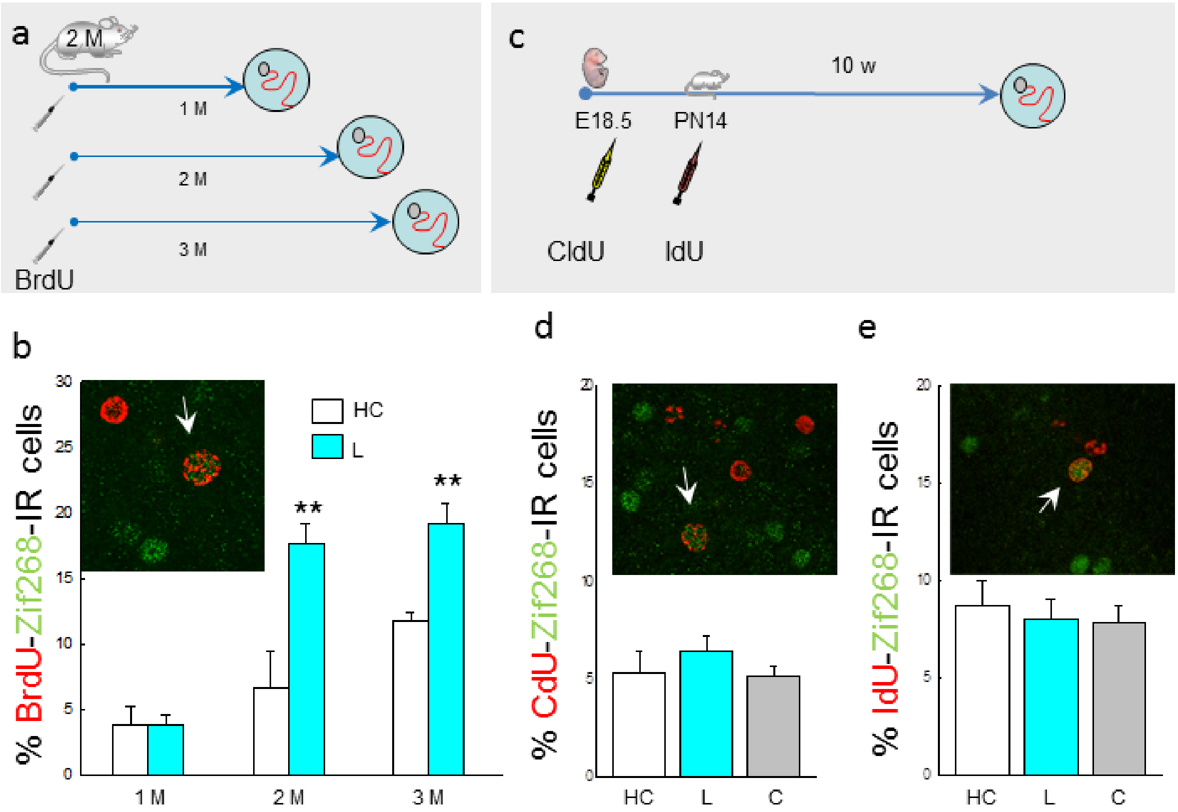


**Figure S1: Mature Adu-DGNs are activated by learning in contrast to DGNs born in embryos or neonates.** (**a**) Experimental design: DGNs born in 2-month-old rats were tagged with BrdU. Rats were tested 1, 2 or 3 months after the injections. (**b**) Illustration and quantitative analysis of the percent of BrdU cells expressing zif268. (**c**) Experimental design: DGNs born in embryos were tagged with CldU (injected to the pregnant mothers) and DGNs in the same animals born at PN14 were labeled with IdU. Ten weeks after the last injection animals were trained in the water maze. (**d**) Illustration and quantification of the percent of CldU cells (in red) expressing zif268 (in green). **(e)** Illustration and quantitative analysis of the percent of IdU cells expressing zif268. HC: home Cage; L: Learning, C: Cued. Arrows indicate double labeled cells. ** $p<0.01$ in comparison to the HC.

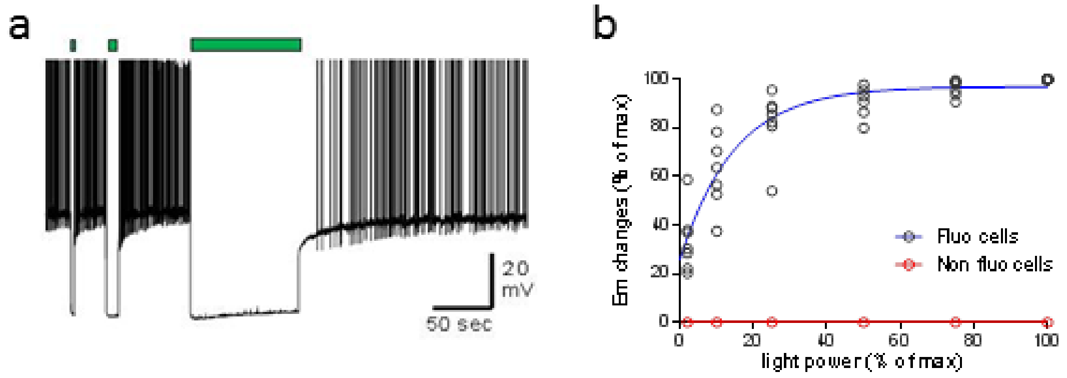


**Figure S2: Whole cell recordings of DGNs in rats infected with RV-Arch-eGFP. (a)** Representative trace showing a whole cell patch-clamp recording under current clamp mode. Positive currents were injected in order to make the cell fire artificially. When the light was turned on (2, 10 and 90 sec, green rectangle), cell potential was instantly reduced and neurons stopped firing. **(b)** Light effect onto neuronal activity, evaluated by monitoring membrane potential (Em), was power-dependent and reached a maximal effect with low power. Importantly, no effect was observed in non-fluorescent cells, in which Em was not changed.

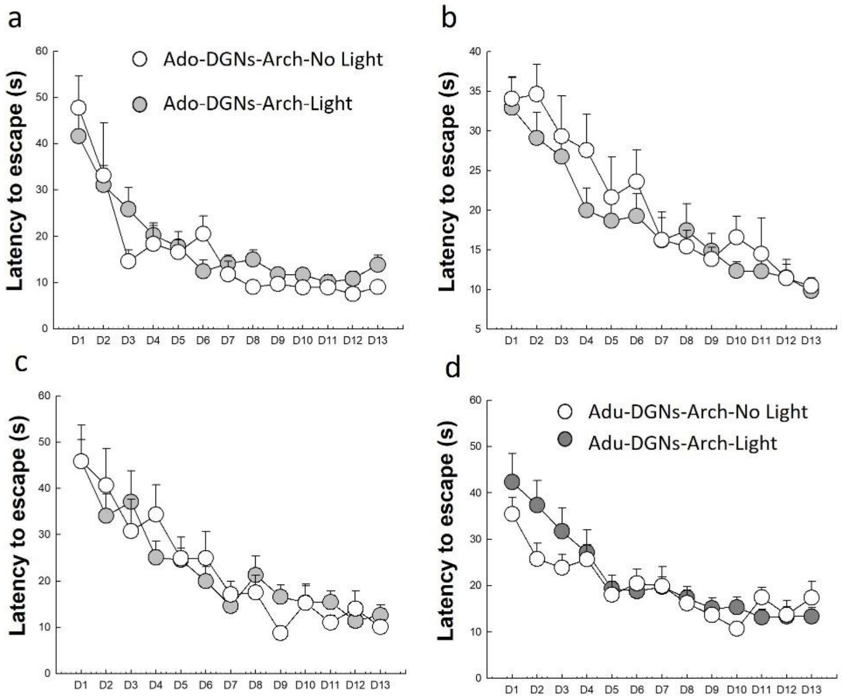


**Figure S3: Optical silencing of DGNs born in adolescent or adult rats does not influence acquisition of spatial memory.** RV-Arch-EGFP were injected bilaterally in the DG of adolescent rats at PN28 (Ado) rats and animals were trained 1 (**a**), 2 (**b**) or 4(**c**) months after the infection. In another experiment, RV-Arch-EGFP were injected bilaterally in the DG of 2-month-old rats (**d**), and animals were trained 6 months later. Rats were trained to find a hidden platform in the water maze with half of the group trained with "Light On" and the other half with "Light Off". In all groups, illumination during training did not influence learning.

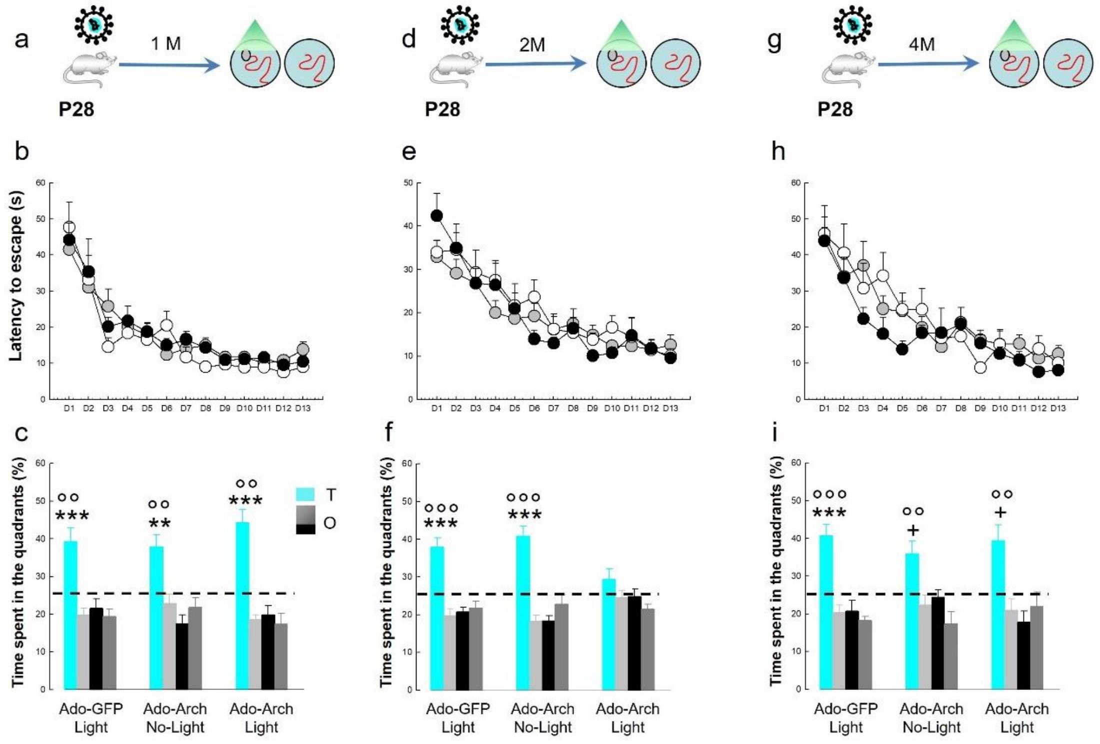


**Figure S4: Light *per se* has no effect in learning nor memory retrieval.** (**a,d,j**) Experimental design: RV-GFP or RV-eGFP-Arch were injected bilaterally in the DG of PN28 rats. (**b,e,k**) Learning curves of GFP-light (black), Arch-No-Light (white), and Arch-Light (grey) groups, where no differences were found. (**c,f,l**) Only silencing 2-month-old Arch-Ado-DGNs during learning impairs the ability of the animals to remember the platform location. $^{+}$ at least at $p<0.05$ and $^{**}p<0.01$. $^{***}p<0.001$ compared to the other quadrants. °° $p<0.01$ and °°° $p<0.001$ compared to chance level. T: target quadrant. O: other quadrants.

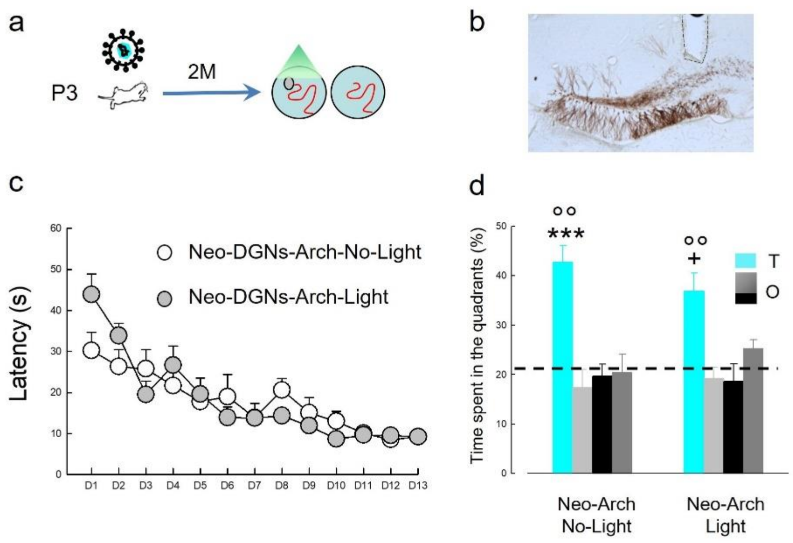


**Figure S5: Optical silencing of DGNs born in neonates does not influence learning nor remembering.** (**a**) Experimental design: RV-Arch-EGFP were injected bilaterally in the DG of neonates at PN3 (Neo). (**b**) Illustration of Neo-DGNs infected with RV-Arch-EGFP; in dashed line, position where the optic fiber was placed. (**c**) Rats were trained to find a hidden platform in the water maze with half of the group trained with “Light On” and the other half with “Light Off”. Illumination during training did not influence learning. (**d**) Silencing 2-month-old Neo-DGNs during learning does not impaired the ability of the animals to remember the platform location. $^{+}$ $p<0.05$ compared to light grey and ***$p<0.001$ compared to the other quadrants. °° $p<0.01$ compared to chance level. T: target quadrant. O: other quadrants.

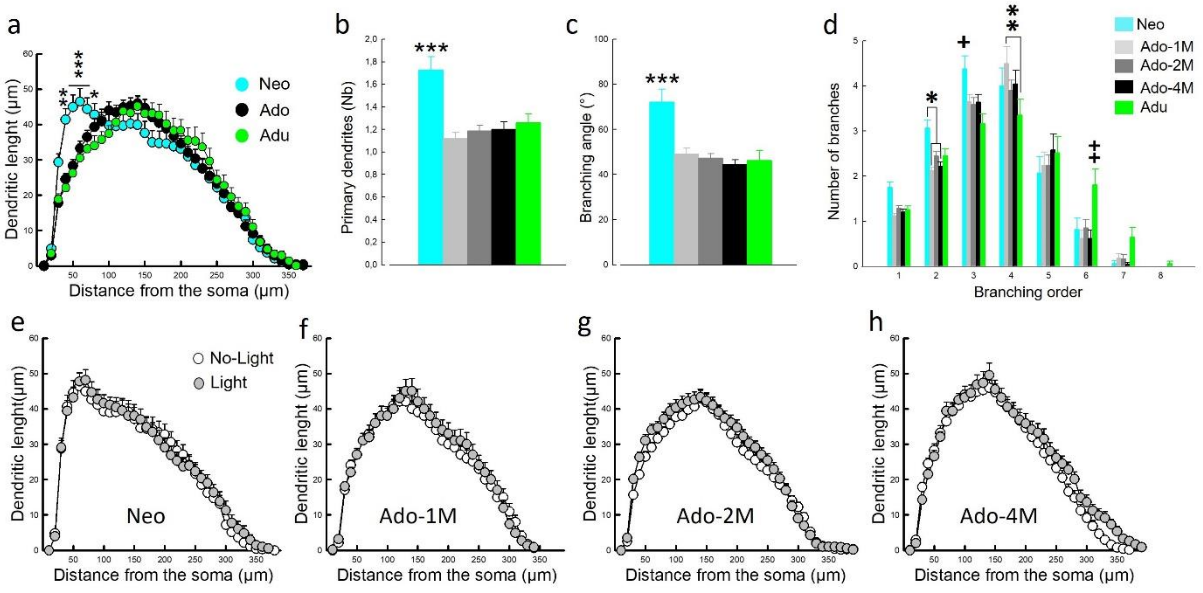


**Figure S6: DGNs generated during development are not affected by the light and present characteristic morphological properties**. Sholl distribution of the three DGNs population analyzed (interaction group x distance : F78, 4095=3.36, p<0.001) shows a different profile in Neo-DGNs (**a**). This population has a higher number of primary dendrites (F4,196=10.34, p<0.0001) (**b**) and broader branching angle (F4,196=11.52, p<0.0001) (**c**) compare with Ado- and Adu-DGNs. Dendritic branching (**d**) shows different profile between the three DGNs population analyzed (interaction group x order : F28,1372=2.57, p<0.0001), but not between the delays in the
